# A pipeline for screening condition-specific enzymes uncovers a function for the alcohol dehydrogenase Bdh2

**DOI:** 10.1101/2025.03.02.641035

**Authors:** Dunya Edilbi, Rosario Valenti, Benjamin Dubreuil, Yeynit Asraf, Sergey Malitsky, Maxim Itkin, Maya Schuldiner

**Affiliations:** Department of Molecular Genetics, Weizmann Institute of Science, Rehovot, 7610001, Israel; Metabolic Profiling Unit, Life Sciences Core Facilities Weizmann Institute of Science, Rehovot, 7610001, Israel

**Keywords:** Enzymes, stress conditions, cell metabolism, metabolites, metabolomics, alcohol dehydrogenase

## Abstract

Cells possess intricate metabolic networks comprised of hundreds of enzymes. Despite extensive research, many of these enzymes remain uncharacterized. Identifying such enzymes is crucial for advancing our understanding of metabolism. However, multiple enzymes are not expressed in standard conditions, making them challenging to study. To overcome this challenge, we created a pipeline for characterizing the expression of condition-specific enzymes in yeast. We assembled a collection of 110 yeast strains, each containing an uncharacterized putative enzyme fused to a fluorophore under the regulation of their own promotor. By subjecting them to 43 diverse growth or stress environments, we identified the biologically relevant conditions for the expression of 19 proteins. We focused on one such putative alcohol dehydrogenase, Bdh2, and functionally characterized it. More broadly, our discovery pipeline lays the foundation for uncovering new condition-specific enzymes. This has implications in cell biology and biotechnology and should expand our understanding of metabolism.

## Introduction

Understanding the intricate workings of the cell necessitates a comprehensive knowledge of all its components. Cellular metabolism consists of the regulated biochemical processes carried out by enzymes organized in metabolic pathways. To preserve homeostasis, a cell adjusts its specific needs by carefully regulating its metabolic pathways (Escoll & Buchrieser, 2023). Therefore, enzyme abundance and activity are highly dependent on the growth environment. While much progress has been made in understanding central metabolic pathways, significant gaps in our knowledge still exist for condition-specific metabolic reactions (Cesur *et al*., 2022). The lack of knowledge about the function of condition-specific enzymes limits our understanding of cell biology as well as the cellular processes involved in development and disease (Yuan *et al*., 2024).

The baker’s yeast, *Saccharomyces cerevisiae* (from here on termed yeast), is a powerful model organism for studying the function of conserved enzymes because it shares most of its proteome with humans (Cohen *et al*., 2022; Kachroo *et al*., 2022) and it is easy to grow and genetically manipulate (Vanderwaeren *et al*., 2022). Indeed, historically, yeast was fundamental in discovering many of the central, and conserved, metabolic pathways, and their enzymes, widening our understanding of metabolism (Caudy *et al*., 2013; Cohen *et al*., 2022; Teng *et al*., 2013).

Even in the well-studied yeast, hundreds of enzymes are still uncharacterized, comprising about a third of the enzyme repertoire (Cohen *et al.,* 2022). While there are many underlying causes, for this, a central one is the narrow grid of growth conditions under which most of the research is performed. Indeed, while “central” metabolism, such as the glycolysis pathway (Lao *et al*., 2022), has been well studied, “peripheral” metabolism such as that activated in unique conditions, remains less explored.

Harmful or extreme environmental conditions can damage vital cellular components leading to cellular stress and pathologies. Therefore, it is important to understand the mechanisms providing tolerance and adaptation to these stresses (Yuan *et al*., 2024). Identifying the enzymes that play a role in such conditions is, therefore, central to better understanding biology under extreme environments. For example, Gpd1 is a key metabolic enzyme that catalyzes the conversion of dihydroxyacetone phosphate (DHAP) to glycerol-3-phosphate in the glycerol biosynthesis pathway. Under osmotic stress, yeast cells upregulate Gpd1 levels to produce more glycerol to balance internal osmotic pressure (Albertyn *et al*., 1994; Pallapati *et al*., 2022). However, many condition-specific enzymes are understudied due to their unique expression patterns, preventing a comprehensive understanding of yeast responses to environmental stress (Lin *et al*., 2022).

To overcome these challenges, we created a pipeline for characterizing condition-specific enzymes. To this end, we assembled a collection of yeast strains, each representing one putative, uncharacterized enzyme that is not expressed in standard growth conditions, fused to a fluorophore, and under the control of its native promotor. We subjected this collection of 110 strains to various stresses and extreme metabolic environments, uncovering conditions inducing the expression for 19 of those genes. Out of those we focused on a poorly characterized alcohol dehydrogenase, Bdh2, whose expression was induced only under extreme starvation conditions. We found that loss of Bdh2 had profound implications on the resistance of yeast to these conditions. We manipulated the expression levels of Bdh2 and obtained metabolic fingerprints of the cells under the applied condition. This allowed us to map the cellular function of Bdh2. These findings enhance our understanding of enzyme roles within the cell, alter our understanding of yeast metabolism during stress, and have implications for biotechnology.

## Results

### Assembling a collection of putative condition-specific enzymes

Uncovering the function of uncharacterized enzymes will deepen our understanding of metabolic networks. To study an enzymatic function, first, we must find the exact conditions where the protein is expressed. To approach this task systematically, we utilized a pre-existing collection (Valenti *et al*., 2025) and from it assembled yeast strains, each genomically encoding one uncharacterized enzyme, not expressed under standard growth conditions, fused on its C terminus to a green fluorescent protein (GFP). This genomic manipulation leaves the enzyme expressed from its natural endogenous promotor (Figure 1, Supplementary Table 1).

**Figure 1:**
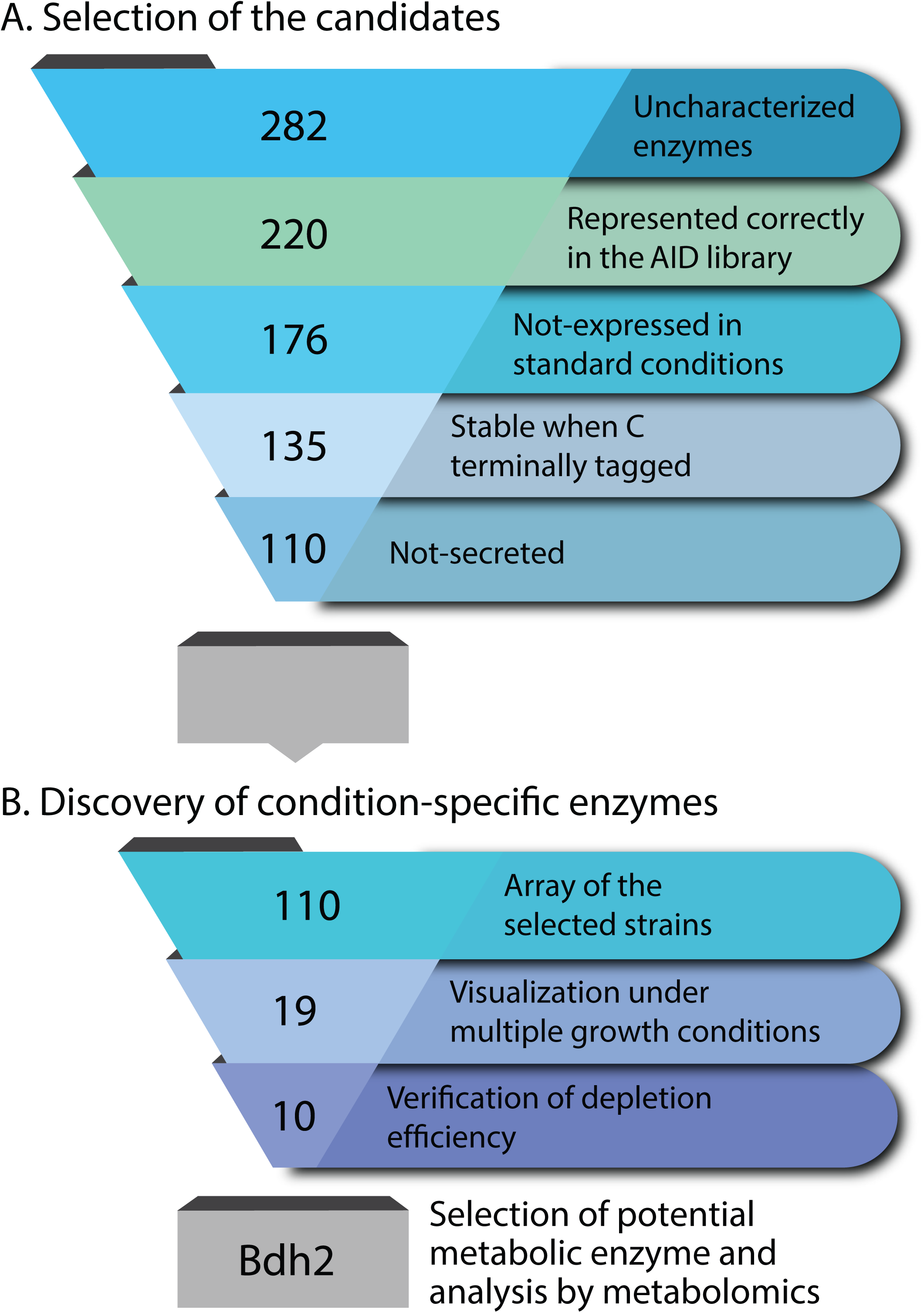
Schematic representation of the selection of candidate enzymes and workflow. A. Schematic representation of the process of selection of the candidates. The uncharacterized enzymes represented correctly in the AID collection and not expressed in standard conditions were our starting point. From these, we removed proteins that we predicted as unstable with a C terminal GFP tag or secreted. This resulted in 110 strains that have a high chance of being condition-specific enzymes. B. Schematic representation of the process of characterizing condition-specific enzymes. We screened 110 strains, each expressing one protein C terminally tagged with GFP in 43 different growth conditions. We verified the capacity of the 19 condition-specific proteins to be depleted by the AID2 system. From the 10 responsive strains, we picked one to characterize in depth.

We started by exploring 282 uncharacterized proteins predicted to have enzymatic function (Cohen *et al*., 2022). Of those uncharacterized candidates, at least 220 were correctly represented in the collection from which we picked the strains (Valenti *et al*., 2025). Out of these, 44 were constitutively expressed. Therefore, we focused on the 176 that did not exhibit a signal in standard growth conditions. We continued to trim our list by removing those strains for which we had an indication that the C terminal tag was not compatible with protein folding or stability, leaving us with 135 strains. To get a sense of the proteins that fall into this category, we searched the LoQAtE database (Breker *et al*., 2013) and retrieved all information on expression when the proteins were C or N terminally tagged and under their native promoter. Those proteins that could be visualized with their native promoter when N but not when C terminally tagged, were considered unstable when C terminally tagged. Next, we removed any enzymes predicted to be secreted since these, even if correctly tagged, expressed, and stable, will be secreted outside the cell, and we would not detect their expression. We defined potentially secreted proteins as having a predicted cleavable signal sequence and no predicted transmembrane domains (Supplementary table 2, Cohen *et al*. 2023). Hence, if an enzyme was represented in our source collection, was not expressed from its native promoter regardless of the tag position, and is not expected to be secreted – then it has a high chance of being regulated for expression under unique conditions. This can be either by transcriptional regulation or by post-transcriptional/translational methods (i.e. by stabilization from degradation). We therefore chose to focus on this group, consisting of 110 candidates (Figure 1).

### Screening diverse conditions for protein expression

To identify putative enzymes expressed under specific growth conditions, we grew the strains in 43 different conditions, including various carbon sources, nutrient changes, growth phases, drugs, and stresses (Table 1). We then imaged the strains using spinning disk fluorescent microscopy (Figure 2A) and identified 19 proteins that were expressed in 12 different conditions. Some proteins were expressed in multiple conditions, but some only in a single condition (Figure 2B). Conversely, some conditions did not affect the expression of any candidate, while others impacted the expression of many (Figure 2B, Supplementary Table 3). Interestingly, although most proteins exhibited cytosolic localization, we found some that localized to specific subcellular locals such as Rix7 and Scw10 to the nucleus, Fmp46 to mitochondria, Fre6 to the vacuole, and Axl2 to budding sites, increasing the ledger of proteins defined in these subcellular compartments (Figure 2A, Supplementary Table 4).

**Table 1:** Conditions screened for unique expression patterns. . A table showing all the conditions used to screen the strains for protein expression, clustered by type of perturbation, in addition to the number of expressed proteins in each condition, in brackets.

| Stressors | Drugs | Carbon sources | Metals |
| --- | --- | --- | --- |
| SD+2DG (6) | Fluconazole (1) | SGlycerol (0) | SD-Fe (0) |
| 5% Ethanol (1) | Amphotericin B (0) | SGalactose (1) | SD-Cu (5) |
| DDW (3) | Terbinafine (0) | SSucrose (0) | SD-Zn (0) |
| SD-aa (5) | Tunicamycin (1) | SRaffinose (3) | SD+EGTA (2) |
| SD+10X aa (6) | CCCP (8) | SOleate (1) | SD-Mg (0) |
| YPD (2) | Rapamycin (0) |  | SD-Mn (3) |
| SD no MSG (3) | Cerulenin (5) |  |  |
|  | DTT (0) |  |  |
| Co-factors | Osmotic stress | Phases | Minerals |
| SD-Thiamin (7) | SD+KCl (0) | Exit Stationery (0) | SD-Ca (2) |
| SD-Inositol (1) | SD+NaCl (0) | Deep stationary (3) | SD+Ca (7) |
| SD-Riboflavin (1) | SD+Sorbitol (0) | Mating (0) | SD-Phosphate (3) |
| SD-Biotin (6) |  | Diploids (0) |  |

**Figure 2:**
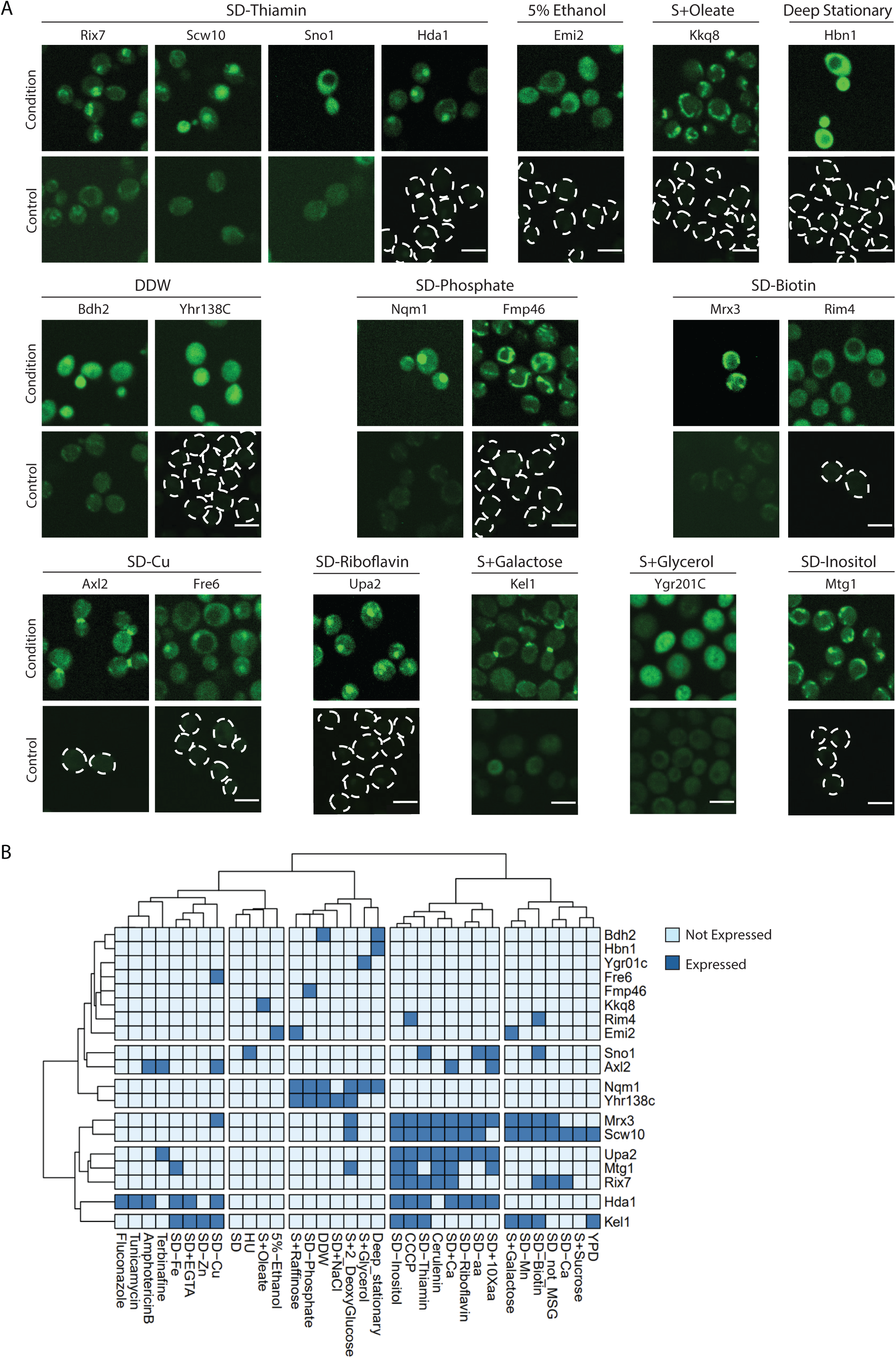
Screening multiple environments uncovers condition-specific proteins. A. Fluorescent images of the strains with the proteins expressed and under which conditions. Shown is also a control of the same strain in normal conditions (SD). Scale bar, 5 µm. B. A cluster plot shows all the strains with the proteins that were expressed and under which conditions.

### Assaying the capacity to deplete proteins under conditions in which they were observed

Our pipeline uncovered a unique expression pattern for 19 putative enzymes, suggesting the conditions in which these enzymes function. One way to study their activity would be to deplete them in these extreme conditions and assay the effects of this manipulation. In anticipation of this, we had already assembled the strains assayed above from a collection that had, in addition to GFP, a tag enabling rapid protein degradation (Valenti *et al*., 2025). This tag encodes for the improved Auxin Inducible Degron system (AID2) (Yesbolatova *et al*. 2020). These strains are therefore unique in allowing us both to track protein expression, abundance, and localization as well as rapidly depleting them by adding a small auxin-like molecule (5-Ph-IAA). However, the activities of the E3 ligase, Tir1, and the ubiquitin-proteasome pathway (the pathway that leads to the depletion of the proteins after 5-Ph-IAA addition) may vary under different conditions, making it unclear whether degradation would be equally efficient across all conditions. To assay the degradation efficiency across the conditions, we took the 19 expressed fusion proteins grown under the conditions in which they were visualized and triggered their degradation. We found that 10 out of the 19 strains were responsive (Figure 3). Since we observed depletion of a control protein under every condition tested (data not shown), we assume there was a specific issue with the remaining nine strains that did not respond to 5-Ph-IAA. These results emphasize the robustness of the AID2 system, allowing reliable protein depletion even under varied metabolic conditions.

**Figure 3:**
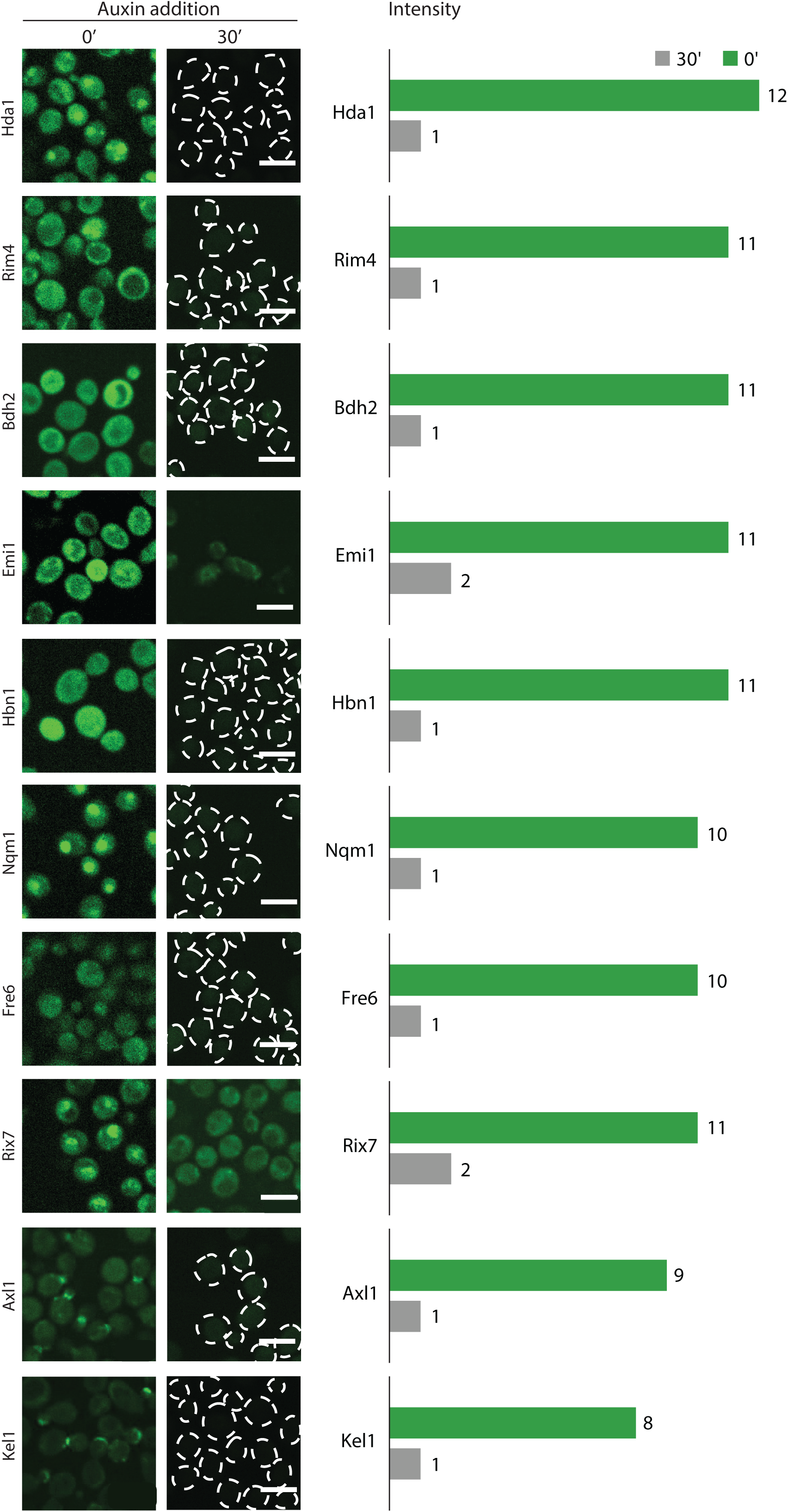
The AID2 system allows the regulation of protein abundance even under unique growth conditions. Fluorescent images showing the strains with the targeted proteins that were responsive to 30 minutes (30’) 5-Ph-IAA addition under the same condition in which they were expressed (left). Scale bar, 5 µm. The fluorescent intensity of each strain, before and after depletion, is represented in the graph (right).

### Bdh2 is a putative alcohol dehydrogenase that has a central role under extreme starvation conditions

Out of the 10 putative enzymes we could deplete using the AID2 system, we focused on an interesting candidate, Bdh2. This protein and its close paralog Bdh1 possess a putative alcohol dehydrogenase domain (Nash *et al*., 2020). Furthermore, the UniProt database (Coudert *et al*., 2023) suggests an EC number (Enzyme Commission number) of 1.1.1.303 for Bdh2, which is butanediol dehydrogenase activity. In support of this, it was demonstrated that even in the absence of Bdh1, Bdh2 affects the degradation of vanillin, a phenolic aldehyde susceptible to reduction to alcohol by an alcohol dehydrogenase (Ishida *et al*., 2016). However, the endogenous substrate or products of Bdh2 have not been clearly demonstrated nor has its physiological role.

Although many alcohol dehydrogenases have been extensively studied, Bdh2’s role has remained unexplored since it is not expressed under standard conditions. Western blot analysis verified that Bdh2 accumulates under two of the screened conditions: deep stationary phase and growth in double distilled water (DDW) (Figure 4A).

**Figure 4:**
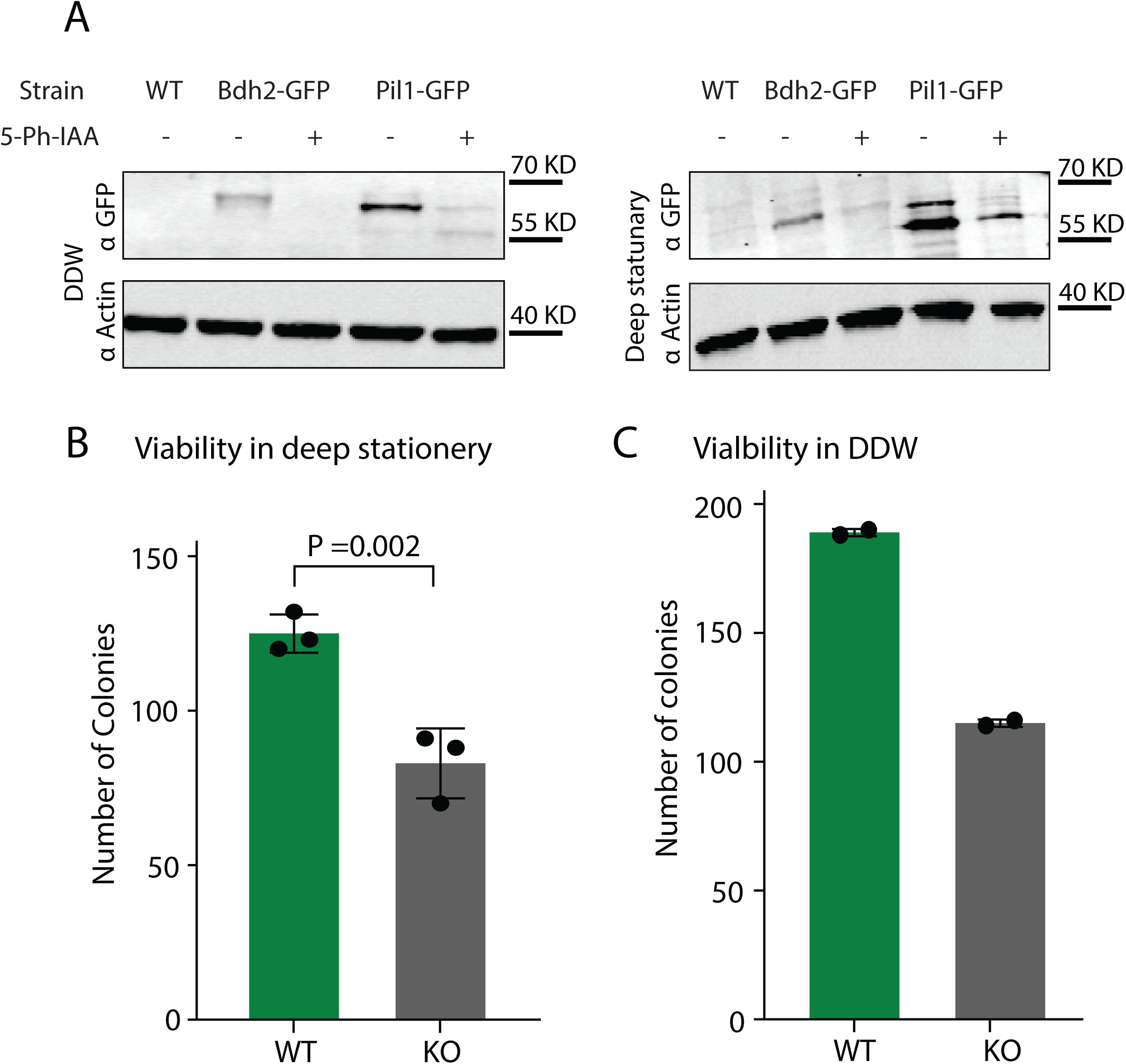
Bdh2 is important in extreme starvation conditions where it is expressed. A. A western blot showing the increased expression of Bdh2-GFP in deep stationery and DDW media. Actin was used as a loading control, and the constitutively active Pil1 protein was used as a positive control. B. A bar graph showing the viability rates for deletion of Bdh2 in deep stationery media relative to control. C. A bar graph showing the viability rates for deletion of Bdh2 in DDW relative to control.

Since both conditions represent an extreme case of nutrient starvation, we tested whether Bdh2 has a role in survival during these stresses. Indeed, loss of Bdh2 resulted in an extreme reduction of viability, manifested by the capacity of colonies to grow after exposure to these conditions (Figure 4B). These findings highlight Bdh2 as a key player in metabolic adaptation to extreme conditions.

### Analysis of metabolic changes induced by altering Bdh2 expression provides clues to its enzymatic activity

To uncover the metabolic effect of altering Bdh2 levels, we assayed the metabolic fingerprint of altering Bdh2 expression. This was done either by depleting Bdh2 using the AID2 system during growth in DDW, a condition in which Bdh2 abundance increased. Alternatively, we either deleted or over-expressed Bdh2 constitutively. We then utilized metabolic profiling (metabolomics) to measure the changes in putative metabolites between the various states (Supplementary Table 5, 6). Since Bdh2 has a predicted alcohol dehydrogenase domain, we focused on changes to either alcohols, aldehydes (precursors to alcohols), or ketones (the products of alcohol dehydrogenation). Indeed, we found four significantly altered putative metabolites from these groups. Two putative metabolites were ketones that were significantly reduced when Bdh2 was depleted and elevated when it was over-expressed: **Phenyl-1,2-propanedione** (Figure 5A) and **Dihydroxyacetone** (Figure 5B). One putative compound was an aromatic alcohol **2-(4-hydroxyphenyl)ethanol,** that was decreased when Bdh2 was over-expressed. In addition, we noticed that during Bdh2 depletion its levels decreased although this was not statistically significant (Figure 5C). The fourth putative compound was an aldehyde **3-(4-Hydroxyphenyl)propionic acid**, that was significantly increased in the over-expression strain and reduced in the depleted strains (Figure 5D). Interestingly, some compounds did not change significantly in the deletion strain but did in the depletion conditions. This, most likely, stems from the extensive metabolic rewiring that cells undertake following a genetic alteration to maintain homeostasis and underlines the importance of using an “on-demand” degradation system. Regardless, the metabolic changes suggest potential reactions that could be carried out by Bdh2 *in vivo* during survival in DDW (Figure 5A, B, C, D).

**Figure 5:**
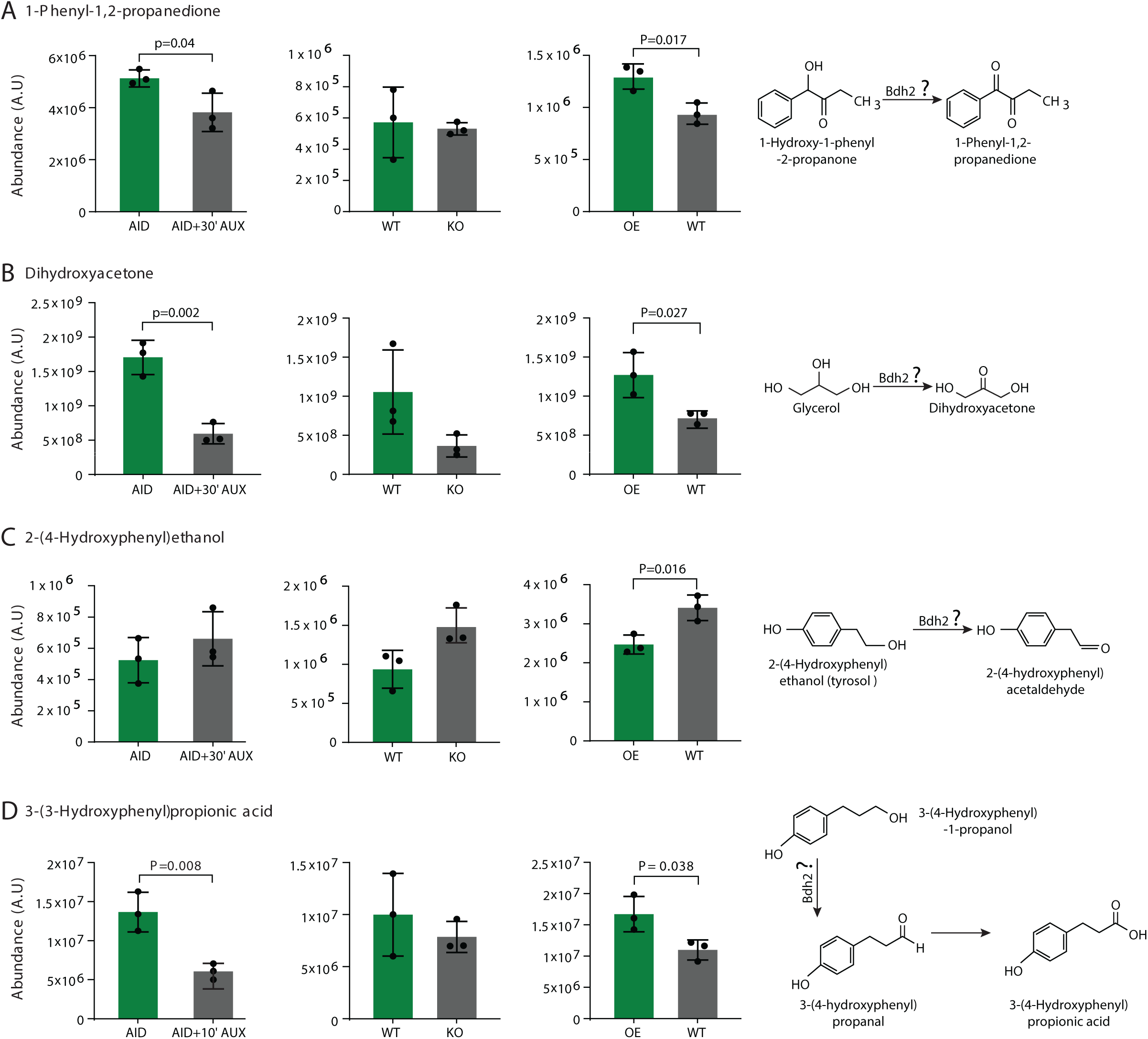
Metabolomics analysis suggests a role for Bdh2, a predicted alcohol dehydrogenase. A. Bar graphs showing the changes in the ketone 1-Phenyl-1,2-propanedione upon depletion, knock out, and overexpression of Bdh2 compared to control, in addition to the proposed reaction in which 1-Phenyl-1,2-propanedione takes place. B. Bar graphs showing the changes in the ketone Dihydroxyacetone upon depletion, knock out, and overexpression of Bdh2 compared to control, in addition to the proposed reaction in which Dihydroxyacetone takes place. C. Bar graphs showing the changes in the aromatic alcohol 2-(4-hydroxyphenyl)ethanol upon depletion, knock out, and overexpression of Bdh2 compared to control, in addition to the proposed reaction in which 2-(4-hydroxyphenyl)ethanol takes place. D. Bar graphs showing the changes in the acid 3-(4-Hydroxyphenyl)propionic acid upon depletion, knock out, and overexpression of Bdh2 compared to control, in addition to the proposed reaction in which 3-(4-Hydroxyphenyl)propionic acid takes place.

Together our metabolic analyses support a role for Bdh2 as an alcohol dehydrogenase whose activity is essential for survival in extreme starvation conditions.

## Discussion

This study aimed to create a conceptual pipeline for discovering functions of uncharacterized enzymes in yeast that are only expressed under specific environmental or stress conditions. Using a mini collection of 110 strains, we identified 19 putative enzymes expressed across 12 distinct conditions, such as nutrient deficiencies, oxidative stress, and exposure to metabolic inhibitors. By applying the AID2 system (Valenti *et al*., 2025), we managed to dynamically deplete 10 of the proteins under the conditions in which they were expressed. We focused on one such putative enzyme, Bdh2, whose expression was induced during growth in deep stationary phase or DDW. We found that loss of Bdh2 in these two extreme starvation conditions dramatically reduces yeast viability. Using metabolomic analysis on strains in which we altered Bdh2 expression, we suggest biological roles for Bdh2.

Despite these significant findings, many enzymes in the library were not visualized even though we assayed 43 different conditions. While several factors may contribute to this, we hypothesize that many of the enzymes are expressed at very low levels, which makes them difficult to detect using fluorescent microscopy. To address these challenges, future studies could involve alternative tagging strategies or explore broader environmental conditions.

We focused on Bdh2 and found that it may have a unique role in DDW and stationary phase – two conditions where nutrients are sparse. One putative metabolite that showed coherent changes was **2-(4-hydroxyphenyl)ethanol**, also known as tyrosol, a fusel alcohol formed as part of the Ehrlich pathway (Ehrlich., 1907). The catabolism of amino acids is a valuable source of nitrogen during starvation and is a multi-enzyme process (Hazelwood *et al*., 2008). However, the specific contributions of each isozyme under various physiological conditions in the Ehrlich pathway are not fully delineated (Hazelwood *et al*., 2008). Further research is needed to verify if Bdh2 could indeed be one of the alcohol dehydrogenases acting on the Ehrlich Pathway. An additional putative metabolite that was affected was **Dihydroxyacetone,** which is derived from glycerol, a byproduct of phospholipid catabolism (Klein *et al*., 2017). In conditions of nutrient scarcity, fats are mobilized to sustain cellular energy requirements, leading to glycerol release, which might subsequently be metabolized by Bdh2 (Kim *et al*., 2023). However, our metabolomics data shows changes in glycerol that are not simple to explain by such a reaction, leaving it open whether this effect is direct or not. Regardless of the specific substrate or product, the chemical reaction performed by Bdh2 is expected to generate NADH, which is essential for both respiratory chain complex activity during respiration as well as maintaining redox balance (Yuan *et al*., 2024). This is important since yeast mostly respire during nutrient-limiting conditions (Tu *et al*., 2007).

In contrast, Bdh1, the paralog of Bdh2, is constitutively expressed, potentially since it serves a housekeeping role in alcohol metabolism under normal growth conditions, continuously contributing to fermentation and energy turnover (Nash *et al*., 2020).

Overall, our study identified putative enzymes that are expressed under specific environmental conditions. It provides a roadmap for future research into the characterization of enzymes that are not expressed under standard laboratory conditions and, therefore, may play a role in peripheral metabolism and stress resistance. Together these promote our efforts to uncover more enzyme functions and should advance our understanding of metabolic networks and their regulation.

## Materials and methods

### Yeast strains

We picked 110 strains of uncharacterized enzymes and 6 positive control strains from the same collection (Valenti *et al*. 2025) and added 4 negative control strains, to generate our mini collection of uncharacterized and unexpressed enzymes. All the strains are listed in Supplementary Table 1. All Primers were designed using the Primers 4 Yeast web tool (Yofe & Schuldiner, 2014) (Supplementary Table 7).

### Yeast growth

Yeast cells were grown on solid media containing 2.2% [w/v] agar (Formedium #AGA0X) or liquid media, at 30°C. Antibiotics: nourseothricin (NAT, WERNER BioAgents “ClonNat” #5.XXX.000) at 0.2 g/l and geneticin (G418, Formedium #G418X) at 0.5 g/l were used for strain maintenance and overnight pre-cultures. Unless stated otherwise, the media used was synthetic minimal media (SD; 0.67% [w/v] yeast nitrogen base (YNB) without amino acids or ammonium sulfate (Formedium #CYN04XX),0.1% L-Glutamic acid monosodium salt, 2% [w/v] glucose), supplemented with amino acid OMM mix (Hanscho *et al*. 2012) (SD (MSG) Complete). For the expression tests and microscopy, the strains were grown in different media conditions (Table 1, Supplementary Table 3).

### Automated high-throughput fluorescence microscopy

For high throughput fluorescence microscopy imaging, the strains were transferred from agar plates into 384-well plates for growth in 100µl condition-specific media (Supplementary Table 3) overnight (ON) at 30°C using a RoToR arrayer (Singer). The ON culture was back diluted into 384-well plates in condition-specific media to an OD_600_ of ∼0.2. After 4h, 50µl from each well was transferred to a glass-bottom 384-well microscopy plate (Azenta Life Sciences) coated with concanavalin A (Sigma-Aldrich). The cultures were allowed to adhere to the bottom of the plate for 20 minutes. After incubation, wells were washed twice with condition-specific media to remove non-adherent cells, all was done in EVO freedom liquid handler (TECAN). The plates were then transferred to an Olympus automated inverted fluorescent microscope system using a robotic swap arm (Peak Analysis & Automation). Cells were imaged at room temperature (RT) in condition-specific media using a 60X air lens (NA 0.9) and with an ORCA-flash 4.0 digital camera (Hamamatsu). Images were recorded with 488nm laser illumination for the GFP channel (excitation filter 488nm, emission filter 525/50nm), with 600ms exposure and 60% laser intensity, mild conditions to minimize photobleaching. Bright-field images were also taken, with 200ms exposure and 100% LED power. Four positions were imaged per well and the software autofocus was used to ensure the cells were imaged in their central plane for proper comparison (scanR version 3.2).

Images displayed in Figures 2 & 3 were acquired at RT using a VisiScope Confocal Cell Explorer system, composed of a Zeiss Yokogawa spinning disk scanning unit (CSU-W1) coupled to an inverted Olympus IX83 microscope with a 60X oil objective (NA 1.4). The excitation wavelength was 488nm, with an exposure time of 600ms and 80% LED intensity). Images were taken by a PCO-Edge sCMOS camera controlled by VisiView 3.2.0 software.

The microscopy images were cropped and attributed with a color gradient for representing the range of fluorescence intensity in ImageJ (Schneider, Rasband, and Eliceiri 2012). The brightness and contrast of images were linearly adjusted so that all GFP images of the same strain (before and after induction) have the same parameters.

### 5-Ph-IAA-induced degradation assays

Protein depletion was assayed under some of the tested conditions (Supplementary Table 3), after 30min of 5µM 5-Ph-IAA (an Auxin analog, Bio Academia: Product code: 30-003-10). Addition of 5µM 5-Ph-IAA was in the lag phase (30min before the back dilution time ends), and the cells were washed with media plus 5µM 5-Ph-IAA for imaging.

### Western blot analysis

Cells were grown ON in rich media (YPD: YPD broth 5% [w/v] Formedium #CCM02XX: 2% [w/v] peptone, 1% [w/v] yeast extract, 2% [w/v] glucose). With selections and used to inoculate 20ml of YPD to OD_600_ 0.05. Cells were incubated with shaking at 30°C until OD_600_ 0.2, where they were split into 2 identical cultures. One culture received 5µM 5-Ph-IAA (30min before the final time point), and the other culture was treated with DMSO as control. Cells were harvested after 3.5h, except for strains in DDW that were incubated for another 4h, and cells grown in deep stationery media that were incubated for 24h, to detect the effect of the stress on protein expression. Cells were harvested by centrifugation followed by flash freezing in liquid nitrogen. Protein extraction, SDS-PAGE, and western blotting were performed as described previously (Eisenberg-Bord *et al*., 2021). Briefly, the cells were lysed with 8M urea-based lysis buffer with protease inhibitors by glass bead-beating. Lysates were denatured by the addition of SDS (final concentration ∼2%) and a 45°C incubation for 15min. Denatured lysates were centrifuged to separate cell debris. Loading buffer containing DTT (final concentration ∼25mM) was added, and samples were incubated at 45°C for 15min. 30μg sample was loaded onto 10% agarose gels and separated with electrophoresis, then transferred onto nitrocellulose membrane using the Trans-Blot Turbo transfer system (Bio-Rad). Membranes were blocked in SEA BLOCK buffer (Thermo Scientific), incubated with primary antibodies (anti-GFP, Abcam, ab290 1:1000 and Anti-actin, Abcam, ab170325 1:5000), washed, and incubated with fluorescent secondary antibodies for 1h (Li-COR, 926-32210, 1:10000 and Abcam, ab216777, 1:10000). After washing, the membranes were imaged on the LI-COR Odyssey Infrared Scanner.

### Viability assays

For the viability essays, the strains were grown ON in standard media. Next, 10mL cultures at an OD_600_ of 0.1 were incubated in triplicates or duplicates for 4h in Deep stationery and DDW and left in the incubator at 30°C. Then 10µL of each culture, at time 0 and after 3 days, was diluted in 1ml of DDW, and 100ul dilution was plated on agar plates with selections. The plates were grown in 30 for two days. After two days the number of colonies from each plate was counted and compared between the depletion of Bdh2 and the control.

### Sample preparation for metabolomics

Starter cultures were prepared by inoculating 3mL of SD complete medium, with selections, with each strain in triplicates and incubating ON at 30°C on a shaker. Then, 50µL of the overnight cultures were used to seed 5mL fresh cultures of SD complete or condition-specific media (without antibiotics) and incubated ON at 30°C on a shaker. Next, 50mL cultures at an OD_600_ of 0.5 were incubated for 4h and 5-Ph-IAA (2µM final concentration) was added to half of the samples followed by a 30min or 10min incubation at 30°C. Samples were harvested by centrifugation (3000 rpm for 3 minutes), washed in 1mL DDW, centrifuged, and after removal of the DDW, the pellets were snap-frozen in liquid nitrogen.

### Metabolite extraction

Extraction and analysis of polar metabolites were performed as previously described (Malitsky *et al*. 2016; Zeng *et al*. 2015), with some modifications: yeast cell pellets were extracted with 1ml of a pre-cooled (−20°C) homogenous methanol-methyl-tert-butyl-ether (MTBE) 1:3 (v/v) mixture. The tubes were vortexed and then sonicated for 30 min in an ice-cold sonication bath (taken for a brief vortex every 10 min). Then, DDW-methanol (3:1, v/v) solution (0.5mL) containing internal standard, C13, and N15 labeled amino acids standard mix (Sigma, 767964), was added to the tubes, followed by centrifugation. The upper organic phase was transferred into a 2mL Eppendorf tube. The polar phase was re-extracted as described above, with 0.5mL of MTBE, and the organic phase was combined with the first extraction. The polar phase was moved to a new Eppendorf, dried for 1.0h by N2 to remove organic solvents, and then lyophilized. Before the injection into the LC-MS, the polar phase sample pellets were dissolved using 150μl DDW-methanol (1:1), centrifuged twice (13,000 rpm) to remove possible precipitants, and transferred to an HPLC vial.

### LC-MS polar metabolite analysis

Metabolic profiling of the polar phase was performed as described (Zheng *et al.,* 2015) with minor modifications described below. Briefly, analysis was performed using Acquity I class UPLC System combined with mass spectrometer Q Exactive Plus Orbitrap™ (Thermo Fisher Scientific), which was operated in a negative ionization mode. The LC separation was done using the SeQuant Zic-pHilic (150mm × 2.1mm) with the SeQuant guard column (20mm × 2.1mm) (Merck). The Mobile phase B - acetonitrile and Mobile phase A - 20mM ammonium carbonate with 0.1% ammonia hydroxide in DDW - acetonitrile (80:20, v/v). The flow rate was kept at 200μl min^−1,^ and the gradient was as follows: 0-2min 75% of B, 14min 25% of B, 18min 25% of B, 19min 75% of B, for 4min.

### Statistical analysis of the metabolic data

For the statistical comparison between metabolite levels before and after protein depletion, a t-test with a two-tailed distribution and a two-sample equal variance (homoscedastic) were used. Presented tabs include normalized data, containing metabolite names, relative abundance, identification levels, and chemical formula, in addition to charts of averages and standard error bars, basic statistical analysis (t-test), Principal component of Analysis (PCA), possible co-eluting compounds (with the same fragmentation, mass, and similar retention time) (Supplementary material 5, 6).

### Image analysis

The images of the cells were detected, segmented and quatified by the software, scanR Analysis (version 3.2). The intensity of GFP fluorescence (Figure 3) was calculated by deducting the mean intensity of all the negative controls of the collection (non-GFP containing cells) from the total intensity of each image.

### Expression heatmap generation and clustering methodology

We analyzed condition-specific expression patterns of 19 putative enzymes across 35 environmental conditions through heatmap visualization and dual clustering approaches. Fluorescence signals were thresh-holded to create a binary presence/absence matrix, where rows represent enzymes and columns represent experimental conditions. A value of 1 is attributed if the enzyme was detected in condition *j* and 0 otherwise. Conditions with no enzyme detection were excluded from the matrix. To cluster enzymes exhibiting similar expression profiles across conditions, we computed the Tanimoto distance on the rows using the default implementation in the *dist()* function from the proxy package in R (Calier *et al*., 2023). This distance relies on counting all types of pairwise matches between rows: 1–1 matches (A), 1-0 matches (B), 0-1 matches (C) and 0–0 matches (D). By comparing the identical matches (A and D) to the total number of pairs, we can determine the similarity coefficient between a pair of enzymes, which can vary between 0 and 1. We obtain the Tanimoto distance (or dissimilarity coefficient) by subtracting the similarity coefficient from 1:

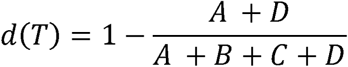

To cluster conditions by similarity of enzymes expressed, we applied a custom version of the Tanimoto distance where we attribute significantly reduced weight for 0-0 matches (D) where both conditions lack expression of the same gene. This weighting scheme reflects the biological consideration that shared enzyme presence provides stronger evidence for condition similarity than mutual absence, given the limited enzyme repertoire relative to the almost infinite universe of conditions.

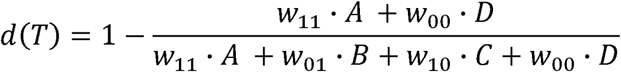

Under the weights w_11_ = w_01_= w_10_=1 and w_00_=0, the formula simplifies to the standard (unweighted) Tanimoto distance for binary data. Finally, for visualization, we used the pheatmap() function from pheatmap package R (Kold., 2018). We performed hierarchical clustering using the hclust() function in R with the “ward.D2” linkage method to iteratively merge clusters while minimizing the total within-cluster variance (Murtagh *et al*., 2014). This agglomerative algorithm optimizes cluster compactness by calculating the increase in variance when merging two clusters and selecting the pair that results in the smallest incremental variance gain.

## Acknowledgements

We thank Prof. Naama Barkai and Dr. Offir Lupo for their helpful input about transcriptional conditions. We thank Yoav Peleg and Shira Albeck for the protein purification. We are grateful to Hanna Vega for the graphical design. We thank Lior Peer and Olga Beresh for critical reading of the manuscript. We thank Dina Daghash for her advice and insight on alcohol and ketone chemistry. This study was supported by the Minerva Foundation, a Chan Zuckerberg Initiative (CZI) grant (2023-331952), and a grant from the Institute for Environmental Sustainability (IES) of the Weizmann Institute of Science. RV was supported by the Alvin, Myra, and David Kaye Memorial Award for excellence in Life Science related research for international students. The robotic system in the Schuldiner lab was purchased through the kind support of the Blythe Brenden-Mann Foundation. MS is an Incumbent of Dr. Gilbert Omenn and Martha Darling Professorial Chair in Molecular Genetics. MI and SM work was supported by the Vera and John Schwartz Family Center for Metabolic Biology.

